# Insights on the taxonomy and ecogenomics of the *Synechococcus* collective

**DOI:** 10.1101/2020.03.20.999532

**Authors:** Vinícius W. Salazar, Cristiane C. Thompson, Diogo A. Tschoeke, Jean Swings, Marta Mattoso, Fabiano L. Thompson

## Abstract

The genus *Synechococcus* (also named *Synechococcus* collective, SC) is a major contributor to global primary productivity. It is found in a wide range of aquatic ecosystems. *Synechococcus* is metabolically diverse, with some lineages thriving in polar and nutrient-rich locations, and other in tropical riverine waters. Although many studies have discussed the ecology and evolution of *Synechococcus*, there is a paucity of knowledge on the taxonomic structure of SC. Only a few studies have addressed the taxonomy of SC, and this issue still remains largely ignored. Our aim was to establish a new classification system for SC. Our analyses included comparing GC% content, genome size, pairwise Average Amino acid Identity (AAI) values, phylogenomics and gene cluster profiles of 170 publicly available SC genomes. All analyses were consistent with the discrimination of 11 genera, from which 2 are newly proposed (*Lacustricoccus* and *Synechospongium*). The new classification is also consistent with the habitat distribution (seawater, freshwater and thermal environments) and reflects the ecological and evolutionary relationships of SC. We provide a practical and consistent classification scheme for the entire *Synechococcus* collective.

## INTRODUCTION

*Synechococcus* was first described by Carl Nägeli in the mid-19th century (Nägeli 1849) and ever since *S. elongatus* has been considered its type species (holotype). *Synechococcus* were regarded mostly as freshwater bacteria related to the *Anacystis* genus (Ihlenfeldt & Gibson, 1975), which is considered a heterotypic synonym for the *Synechococcus* genus. Species later described as *Synechococcus* were also found in thermal springs and microbial mats (Copeland, 1936, Inman, 1940). With the subsequent discovery of marine *Synechococcus* (Waterbury et al. 1979), which were classified as such based on the defining characters of cyanobacteria, described by Stanier (1971), the genus aggregated organisms with distinct ecological and physiological characteristics. The first analysis of the complete genome of a marine *Synechococcus* (Palenik et al. 2003) already displayed several differences to their freshwater counterparts, such as nickel- and cobalt- (as opposed to iron) based enzymes, reduced regulatory mechanisms and motility mechanisms.

Cyanobacteria of the genus *Synechococcus* are of vital importance, contributing to aquatic ecosystems at a planetary scale (Zwirglmaier et al. 2008, Huang et al. 2012). Along with the closely related *Prochlorococcus*, it is estimated that these organisms are responsible for at least one quarter of global primary productivity (Flombaum et al. 2013), therefore being crucial to the regulation of all of Earth’s ecosystems (Bertilsson et al. 2003). Both of these taxa are globally abundant, but while *Prochlorococcus* is found in a more restricted latitudinal range, *Synechococcus* is more widely distributed, being found in freshwater ecosystems, hot spring microbial mats, polar regions, and nutrient-rich waters (Farrant et al. 2016, Sohm et al. 2016, Lee et al. 2019). This demonstrates the metabolic diversity of *Synechococcus*, which has served as a model organism for biotechnological applications (Hendry et al. 2016). Genomic studies deepened our understanding of the unique adaptions of different lineages in the group, regarding their light utilization (Six et al. 2007), nutrient and metal uptake (Palenik et al. 2006) and motility strategies (Dufresne et al. 2008). By analysing the composition of *Synechococcus* genomes, Dufresne and colleagues (2008) identified two distinct lifestyles in marine *Synechococcus* lineages, corresponding to coastal or open ocean habitats, and although there might be an overlap in geographical distribution, niche partitioning is affected by the presence and absence of genes. These insights were mostly restricted to marine *Synechococcus* genomes, and by then, freshwater strains still had their taxonomy status relatively poorly characterized. With these early genomic studies, clear separations started to show between the freshwater type species *Synechococcus elongatus* PCC 6301 and marine lineages such as WH8102 and WH8109. Gene sequences identified as Synechospongium appear in numerous ecological studies as a major component of different sponge species (Erwin & Tacker, 2008). However, this genus has not been formally described, having an uncertain taxonomic position. Despite remarkable ecological and physiological differences within the *Synechococcus* and the successful identification of distinct genomic clades (Ahlgren & Rocap 2012, Mazard et al. 2012, Farrant et al 2016, Sohm et al 2016), the taxonomy of the Synechococcus collective (SC) remained largely unresolved.

A first attempt to unlock the taxonomy of SC was performed by Coutinho et al (2016ab). They compared 24 *Synechococcus* genomes and i. proposed the creation of the new genus *Parasynechococcus* to encompass the marine lineages and ii. described 15 new species (Coutinho et al. 2016b). The description of these new species was attributed to the genetic diversity within these genomes, approaching the problem of classifying all of them under the same name (an issue previously raised by Shih et al. 2013). The new nomenclature also highlighted the genetic difference between marine *Parasynechococcus* and freshwater *Synechococcus*. Walter et al (2017) further elucidates this difference and propose 12 genera for the SC. However, the limited number genomes examined in this previous study hampered a more fine-grained taxonomic analysis of the *Synechococcus* collective.

The present work performs a comprehensive genomic taxonomy analyses using 170 presently available genomes. By combining several genome-level analysis (GC% content, genome size, AAI, phylogenetic reconstruction, gene cluster profiling), we propose splitting the *Synechococcus* collective into 11 clearly separated genera, including two new genera (*Lacustricoccus* and *Synechospongium*). Genus level definition of prokaryotic organisms has been based on the use of AAI (Konstantinidis & Tiedje 2005, Thompson et al. 2013). Modified versions of AAI have also been employed in defining genus level boundaries (Qin et al. 2014) and evolutionary rates across taxonomic ranks (Hugenholtz et al 2016, Parks et al 2018). Therefore, genera were broadly defined based on an AAI cutoff and supported by further genomic analysis, such as the phylogenomic trees, required to confirm genus level definitions (Chun et al. 2018). Based on the presently available data of *Synechococcus* genomes, we propose a new genome-based taxonomy for the group, splitting the *Synechococcus* collective into 10 clearly separated genera, and the creation of two new genera.

## METHODS

### Data acquisition and processing

All *Synechococcus* genomes (n=229) were downloaded from NCBI Assembly database (Kitts et al. 2015) in February 2020 using the Python package “NCBI Genome Download” (https://github.com/kblin/ncbi-genome-download) and querying for the genus “*Synechococcus*”. The metadata table with NCBI Entrez data generated by the package was used as a template for the metadata master table (Table S1). To ensure a standardized treatment of each genome data, instead of using the preexisting files from the assembly directories available at NCBI, only assembly files (containing complete chromosomes, scaffolds, or contigs) were used for analysis.

### Quality assurance

To infer the completeness of each genome, we used CheckM v1.0.12 (Parks et al. 2015) with the “taxonomy_wf” workflow and default settings. The workflow is composed of three steps: i) “taxon_set”, where a taxonomic-specific marker gene set is generated from reference genomes of the selected taxon (in this case, the genus *Synechococcus*), ii) “analyse”, where the marker genes are identified in the genomes, and iii) “qa”, where genomes are assessed for contamination and completeness based on the presence/absence of the marker genes. CheckM results were then parsed with the Pandas v0.25.1 package (McKinney 2011) in a Jupyter Notebook (Ragan-Kelley et al. 2014). Results for completeness and contamination were then added to the master metadata table (Table S1). For all further analyses, we only used genomes with at least 50% completeness and less than 10% contamination as inferred by CheckM. We also removed 9 genomes that did not bin with any other genomes at a 70% AAI cutoff. Thus, 50 “low quality” and 9 “singleton” genomes were discarded, leaving 170 genomes for downstream analyses.

### GC content and genome size

GC content and genome size statistics were calculated from contigs files downloaded from NCBI using Python functions and are displayed in the metadata table (Table S1). The data was aggregated with Pandas to produce the values in Figure 1 and Table 1. For plotting, the libraries Matplotlib (Hunter, 2007) and Seaborn (Waskom, 2018) were used.

**Table 1:** Genera of the *Synechococcus* collective. In total eleven genera, from which two are proposed in the present study (*Lacustricoccus* and *Synechospongium*). Type genomes were chosen based on specific criteria (see Methods section - Description criteria). Additional information for all genomes can be found in Table S1. GC% and genome size (Mbp) values are shown for means ± standard deviation.

| Genus | #<br>genomes | #<br>species* | Type Genome | NCBI name | Lifestyle | GC content<br>(%) | Size (Mbps) |
| --- | --- | --- | --- | --- | --- | --- | --- |
| *Parasynechococcus* | 47 | 22 | *Parasynechococcus africanus*<br>CC9605 | *Synechococcus* sp. | Marine<br>(oceanic) | 58.14 ± 3.02 | 1.96 ± 0.46 |
| *Pseudosynechococcus* | 41 | 21 | *Pseudosynechococcus*<br>*subtropicalis* WH 7805 | *Synechococcus* sp. | Marine<br>(oceanic) | 56.43 ± 3.19 | 2.22 ± 0.48 |
| *Synechospongium*<br>gen. nov. | 28 | 7 | *Synechospongium spongium*<br>15L | Candidatus *Synechococcus*<br>*spongium* | Symbiont | 61.56 ± 1.14 | 1.86 ± 0.28 |
| *Enugrolinea* | 12 | 3 | *Enugrolinea euryhalinus* PCC<br>7002 | *Synechococcus* sp. | Freshwater | 49.26 ± 0.1 | 3.33 ± 0.11 |
| *Regnicoccus* | 9 | 7 | *Regnicoccus antarcticus* WH<br>5701 | *Synechococcus* sp. | Marine<br>(coastal) | 65.36 ± 2.46 | 2.79 ± 0.51 |
| *Inmanicoccus* | 8 | 5 | *Inmanicoccus mediterranei*<br>RCC307 | *Synechococcus* sp. | Marine<br>(coastal) | 61.04 ± 1.55 | 1.78 ± 0.27 |
| *Leptococcus* | 8 | 2 | *Leptococcus yellowstonii* JA-3-3Ab | *Synechococcus* sp. | Thermophilic | 56.34 ± 2.74 | 3.06 ± 0.1 |
| *Thermosynechococcus* | 6 | 5 | *Thermosynechococcus*<br>*elongatus* BP-1 | *Thermosynechococcus*<br>*elongatus* | Thermophilic | 53.65 ± 0.27 | 2.61 ± 0.06 |
| *Synechococcus* | 5 | 2 | *Synechococcus elongatus* PCC<br>6301 | *Synechococcus elongatus* | Freshwater | 55.27 ± 0.25 | 2.75 ± 0.08 |
| *Lacustricoccus* gen.<br>nov. | 3 | 2 | *Lacustricoccus lacustris* TousA | *Synechococcus lacustris* | Brackish | 51.81 ± 0.72 | 1.98 ± 0.62 |
| *Magnicoccus* | 3 | 2 | *Magnicoccus sudatlanticus*<br>CB0101 | *Synechococcus* sp. | Marine<br>(coastal) | 63.43 ± 0.56 | 2.53 ± 0.23 |

\* Several genomes were added to species that were previously defined (in Walter et al 2017) by a single genome. These include, but are not limited to: *Pseudosynechococcus sudipacificus*, *Parasynechococcus marenigrum*, *Inmanicoccus mediterranei*, and, most notably, *Enugrolinea euryhalinus* and *Leptococcus yellowstonii*, respectively with 8 and 7 genomes. In addition to the support of previous species groups, our analysis also expands upon existing genera by proposing new, robust species groups inside of them, specially in *Parasynechococcus*, with 3 new species (with type genomes N32, CC9616, and KORDI-49), containing a total of 16 genomes, and *Pseudosynechococcus*, with 5 new species (with type genomes MITS9504, MITS9508, AG-673-F03, BS55D, and UW105), and a total of 20 genomes. Type species for each species group are noted by a “T” character besides their name (Figure 2). The discovery of these new species can be attributed to a surge of newly available *Synechococcus* high quality whole genome data, obtained mainly from single-cell sequencing (Berube et al. 2018, Kent et al. 2019).

**Figure 1:**
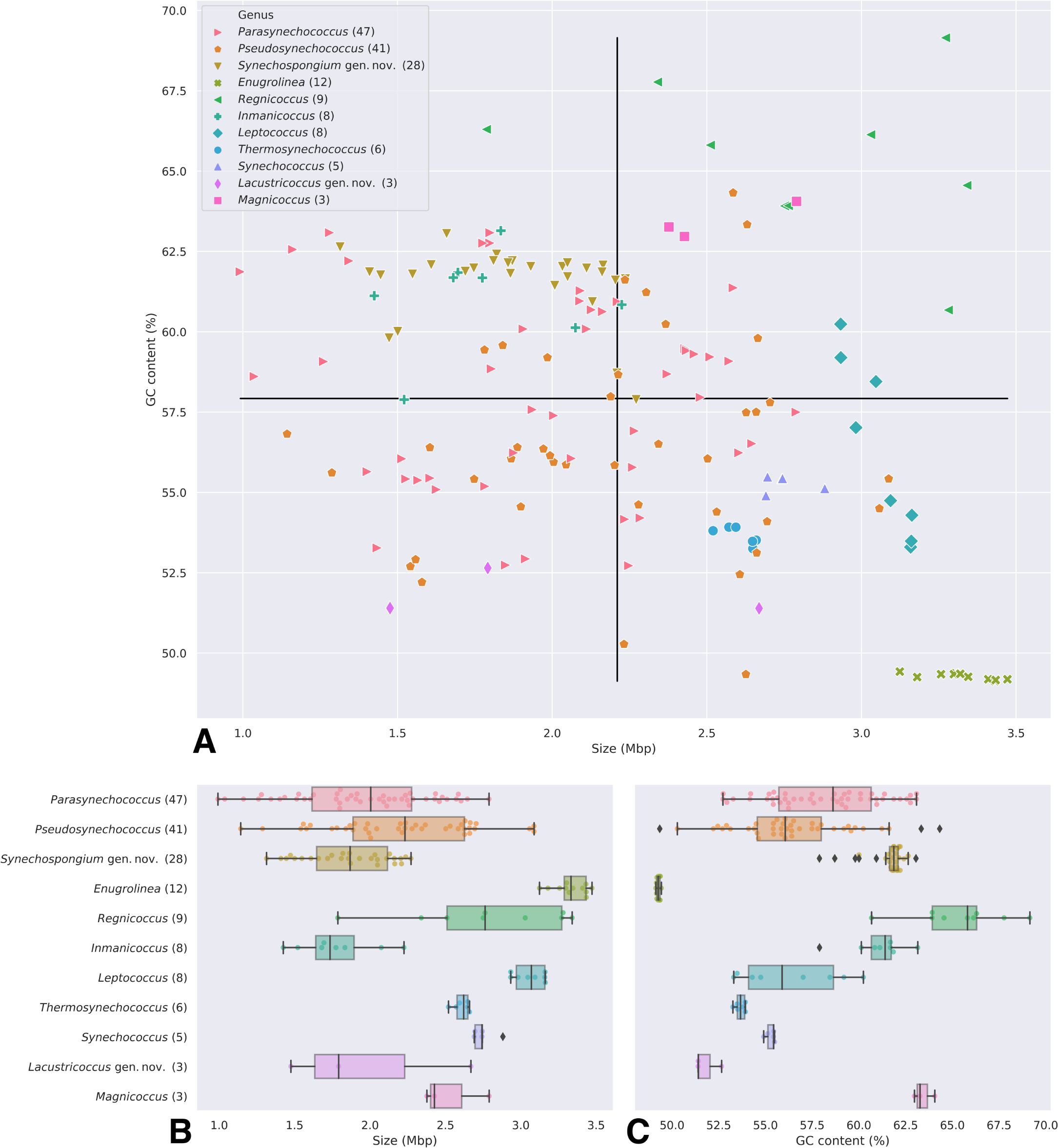
GC content and genome size charts. **A.** Scatter plot of GC content and genome size (in megabases). Black lines indicate the median for all genomes. Genera with lower genetic variability (as shown in the AAI dendrogram) cluster together in small GC/size ranges (with the exception of *Synechospongium* gen. nov.). The genera with most genomes (*Parasynechococcus* and *Pseudosynechococcus*) display a variable GC/size range but still there are no outliers. **B** and **C.** Box plots of genome size (**B**) and GC content (**C**) for each genus. Outliers are shown in diamond shapes. Error bars represent the 1st and 4th quartiles, boxes represent 2nd and 3rd quartiles and the median.

### AAI analysis

Comparative Average Amino acid Identity (AAI) analysis was carried out with the CompareM package (https://github.com/dparks1134/CompareM) v0.0.23. To do so, we ran CompareM’s “aai_wf”, which utilizes protein coding sequences (CDS) predicted with Prodigal (Hyatt et al. 2007), performs all-vs-all reciprocal sequence similarity search with Diamond (Buchfink et al. 2014) and computes pairwise AAI values based on the orthologous fraction shared between genes of the two genomes. The command was run on default settings, with parameters for defining homology being >30% sequence similarity and >70% alignment length. The output table from the AAI analysis was then imported into a Jupyter Notebook a symmetrical distance table was constructed using Pandas v0.25.1. This table is the transformed into a one-dimensional condensed distance matrix using the “squareform()” function from the SciPy library (Jones et al. 2001), “spatial” package. This resulting matrix is subjected to clustering with the “linkage()” function (SciPy library, “cluster” package) with the “method=‘complete’”, “metric=‘cityblock’” and “optimal_ordering=True” parameters. A more in-depth explanation of these parameters can be found in the SciPy documents page (https://docs.scipy.org/doc/scipy/reference/index.html). The resulting array is used as input into a customized function based on SciPy’s “dendrogram()” function.

For our analysis, we performed a hierarchical clustering of pairwise AAI values between all 139 genomes, defining a >70% cutoff for genera (Figure 2). This cutoff is empirically defined by previous studies (Thompson et al. 2013, Rodriguez & Konstantinidis 2014, Qin et al. 2014). Genomes which didn’t cluster with any other genomes based on this criterium were removed from downstream analyses.

**Figure 2:**
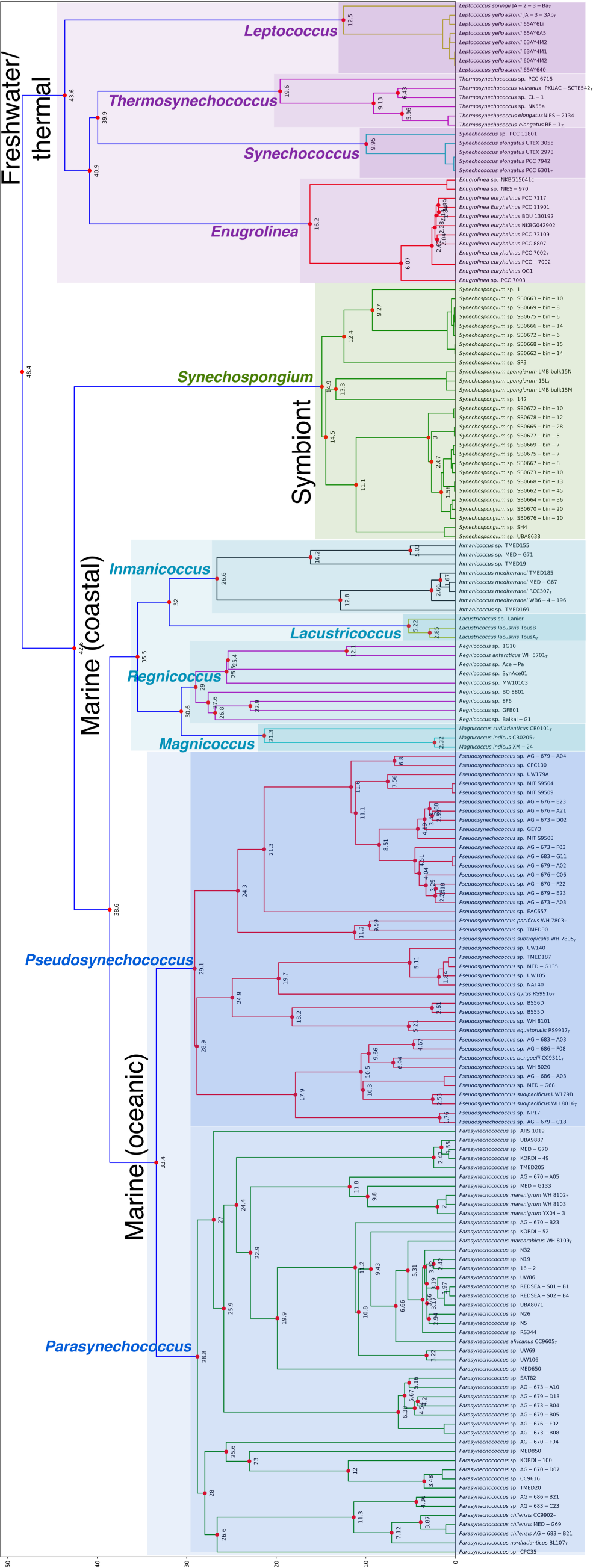
Hierarchical clustering of pairwise AAI values between all *Synechococcus* genomes. New proposed genera are shown within a >70% AAI cutoff. Dotted values show AAI ‘dissimilarity’ values (e.g. 100 minus the AAI value for the pairwise comparison). Dotted values < 1.5 were omitted. Species were defined at a >5% AAI cutoff (Thompson et al. 2013). Type genomes for each SLB are signaled with a “T” character next to the strain name, based on defined criteria (see Methods section). New species were left named as “sp.”. Economic groups are labeled and highlighted in either blue, cyan, green, or purple.

Names for each genera were maintained the same as in Walter et al (2017). An exception to that are the newly-named *Synechospongium* gen. nov. and *Lacustricoccus* gen. nov. Species were defined at a >5% AAI cutoff (based on Thompson et al. 2013). New species were left unnamed. To define a type genome for each species, we used the following criteria, in order of priority: Whether the genome had already been used as a type genome; Genome completeness; Genome release date; Genome source (with a preference for single-cell, then isolate, then metagenome-augmented genomes).

### Phylogenetic trees

To build the phylogenetic trees, we used the GToTree package (Lee, 2019) with default parameters. Two trees were generated, the first (Figure 3, panel A) using 251 Cyanobacteria marker genes and the second (Figure 3, panel B) using 74 Bacteria marker genes. The input dataset consisted of the 170 quality-filtered *Synechococcus* genomes with the addition of a *Prochlorococcus marinus* genome (strain CCMP1375, Genbank accession GCA_000007925.1) to serve as the root for each tree. The genomes were searched against a Hidden Markov Model of the marker genes using HMMER3 (Eddy, 2011). From the 171 genomes, 162 and 160 genomes were respectively retained in the first and second tree after GToTree’s default settings quality control. A concatenated protein alignment from the marker genes was constructed using Muscle (Edgar, 2004) and subsequently trimmed using TrimAl (Capella-Gutiérrez et al. 2009). The alignment was then used to construct a tree using Fast Tree 2 (Price et al. 2010) with default parameters and the pairwise distance matrix using MEGA 6.0 (Tamura, 2013). All processing was done with GNU Parallel (Tange 2018). Trees were rendered using ETE 3 (Huerta-Cepas et al. 2016).

**Figure 3:**
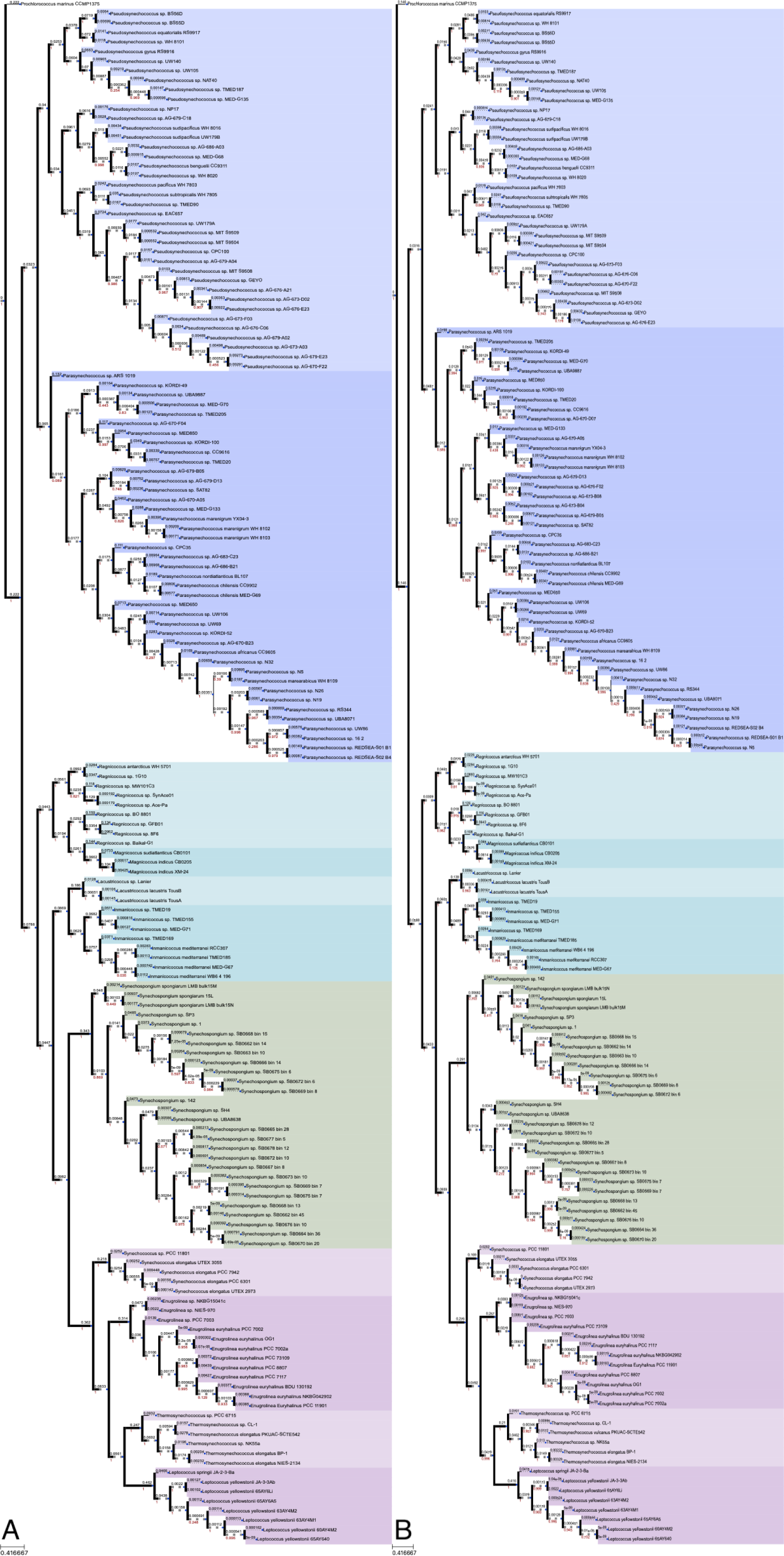
Phylogenetic trees of *Synechococcus-*related genera. Built from the concatenated protein alignment of A) 251 cyanobacterial marker genes and B) 74 bacterial marker genes. *Prochlorococcus marinus* CCMP 1375 is rooted as the outgroup. Red values show branch support and black values show substitutions per site. Ecogenomic groups are highlighted in either blue (Marine/oceanic), cyan (Marine/coastal), green (Symbiont), or purple (Freshwater/thermal).

### CyCOG profiles and *k*-means analysis

Cyanobacterial Clusters of Orthologous Groups profiles were determined by aligning the proteome profiles predicted with Prodigal (see the “AAI analysis” section above) against the NCBI COG database (Galperin et al. 2014) using Diamond in using the parameters ‘evalue=10e-6’ and ‘max_target_alignments=1’. The resulting hits table was filtered against the CyCOG database (Berube et al. 2018), preserving only COGs from cyanobacterial-related genomes. To minimize false negatives gene occurrences, stricter constraints on genome quality were used, and only genomes with at least 95% completeness (as estimated by CheckM) were kept in the CyCOG table. The resulting table (Table S2) was converted to binary form (1 if a CyCOG product was present in a genome and 0 if it was not) and used to plot Figure 4 (CyCOG profiles).

**Figure 4:**
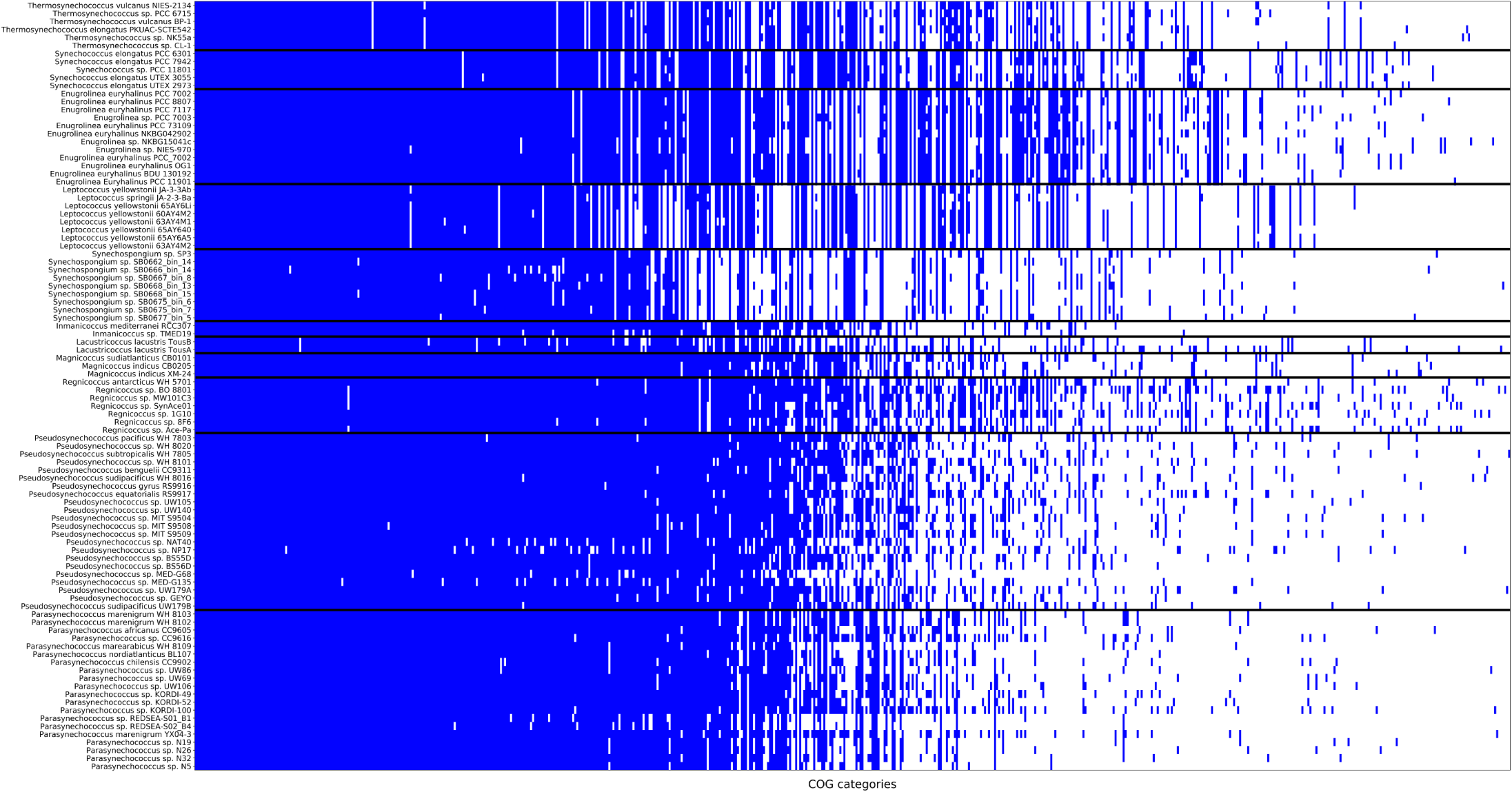
Presence/absence of CyCOG products. Blue bars represent presence of a CyCOG product and white bars its absence for each genome. Different genera are separated by black bars. The data used to generate this figure is in Table S2.

*K*-means analyses were conducted with the implementation available in the SciPy cluster package using the resulting CyCOG table. Values used for *k* were 2, 3, and 4 and the resulting clusters are displayed in Table 2.

**Table 2:** *k*-means groups of CyCOG products. Using the CyCOG presence/absence table, genomes for each genus were clustered using the *k*-means algorithm with *k* values of 2, 3 and 4. All genomes within a genus fell into the same group, therefore it was possible to depict rows as genera instead of individual genomes. As the *k* values increases, it is possible to identify divides within the genera that correspond to ecogenomic groups.

| Genus | 2-means | 3-means | 4-means |
| --- | --- | --- | --- |
| *Leptococcus* | Freshwater/Thermal | Freshwater/Thermal | Thermal |
| *Thermosynechococcus* | Freshwater/Thermal | Freshwater/Thermal | Thermal |
| *Synechococcus* | Freshwater/Thermal | Freshwater/Thermal | Freshwater |
| *Enugrolinea* | Freshwater/Thermal | Freshwater/Thermal | Freshwater |
| *Synechospongium* | Seawater | Symbiont | Symbiont |
| *Regnicoccus* | Seawater | Seawater | Seawater |
| *Pseudosynechococcus* | Seawater | Seawater | Seawater |
| *Parasynechococcus* | Seawater | Seawater | Seawater |
| *Magnicoccus* | Seawater | Seawater | Seawater |
| *Lacustricoccus* | Seawater | Seawater | Seawater |
| *Inmanicoccus* | Seawater | Seawater | Seawater |

### Data and code availability

Whole genome data can be downloaded directly from NCBI Assembly database using the accession codes available in Table S1, in the “assembly_accession” column. We recommend using the above cited “NCBI Genome Download” package to facilitate this. Data generated from CompareM and GToTree and code used for the analysis (in the format of Jupyter notebooks) are available in the following GitHub repository: https://github.com/vinisalazar/SynechococcusGT. Users are encouraged to recreate and examine the figures using Jupyter and the available data. The repository’s “Issues” tab may be used for any further data and/or code requests.

## RESULTS & DISCUSSION

### *Synechococcus* collective GC% content and genome size

Genomic diversity within the *Synechococcus* collective (SC) was observed at several scales, including GC% content and genome size (bp). The sheer span of these two features between genera of the SC indicates marked differences between them. The genome size varies from 0.99 to 3.47 megabase pairs (Mbps), and GC content varies from 49.12% to 69.2% (Figure 1a). However, when the SC is split into several genera, these GC content and genome size values become more consistent (Figure 1bc; Table 1) and closer to proposed ranges for taxonomic grouping (Meier-Kolthoff et al. 2014). Genetically homogeneous genera, such as *Enugrolinea, Synechococcus* and *Leptococcus* form clusters of very low variability in GC content and genome size (Figure 1a). Interestingly, the variability is not so low in the new genera *Synechospongium* (57.89% to 63.05% GC content and 1.31 to 2.27 Mbp) and *Lacustricoccus* (51.9% to 52.6% GC content and 1.47 to 2.67 Mbp).

### Delimitation SC genera by Average Amino acid Identity (AAI)

The AAI analyses discriminated 11 genera (Figure 2). Genomes sharing >70% AAI were grouped into genera. Certain genera (e.g. *Lacustricoccus* and *Synechococcus*) are homogeneous, having at maximum 9.9% AAI difference. Meanwhile other genera (e.g. *Pseudosynechococcus* and *Parasynechococcus*) are very heterogeneous, having up to 29.1% AAI variation. Heterogeneous genera are mostly marine lineages, and display the highest number of genomes (47 and 41, respectively) (Table 1). They are considered oceanic generalists, living in both low and high temperature environments (Walter et al. 2017). In contrast, the freshwater *Lacustricoccus* (previously *Synechococcus lacustris*; Cabello-Yevez et al. 2017, 2018), the thermophilic *Leptococcus*, isolated from Yellowstone hot springs (Becraft et al. 2011), and the *Synechospongium* gen nov. (previously Candidatus *Synechococcus spongiarum*), a symbiont to marine sponges (Usher et al. 2004, Erwin & Thacker 2008, Slaby & Hentschel 2017), appear all to have a more cohesive genome structure at the genus level. The genome previously classified as *Synechococcus lividus* PCC 6715, considered a thermophilic *Synechococcus*, was reclassified as the previously described genus *Thermosynechococcus* (Nakamura et al. 2002), thus enforcing the need to classify novel or earlier *Synechococcus* genomes into a new taxonomic framework. The AAI dendrogram also illustrates the difference between the major ecogenomic groups, which include: Marine/oceanic (*Parasynechococcus* and *Pseudosynechococcus*), Marine/coastal (*Magnicoccus, Regnicoccus, Lacustricoccus* and *Inmanicoccus*), Symbiont (*Synechospongium*), and freshwater/thermal (*Synechococcus* and *Enugrolinea* as freshwater representatives and *Thermosynechococcus* and *Leptococcus* as thermal representatives). The terms “Marine/oceanic” and “Marine/coastal” can also respectively be exchanged “high temperature/low nutrient” and “low temperature/high nutrient” environments.

### Phylogenomic structure of the SC

Genera delimited by AAI analyses were also found by phylogenetic analyses (Figure 3). Both the 251 cyanobacterial marker gene tree and the 74 bacterial marker genes tree depict the eleven genera observed in the AAI dendrogram. The trees support the same groups discriminated in the AAI figure. However, the AAI was superior to discriminate the closely related genera *Magnicoccus* and *Regnicoccus*. These genera group together in both phylogenetic trees, but group separately in the AAI dendrogram (Figure 2). Despite sharing similar ecological characteristics, being sourced from coastal, estuarine-influenced waters, *Magnicoccus* and *Regnicoccus* have distinct GC% and genome size, reinforcing their status as separated genera. The two newly proposed genera (*Lacustricoccus* and *Synechospongium*) form monophyletic branches in both phylogenetic reconstructions, giving strong support for our proposal to formally create these new genera.

### CyCOG profiles and *k*-means analyses

Distinct profiles of Cyanobacterial Clusters of Orthologous Groups (CyCOGs) could be observed for each genus (Figure 4). It is possible to observe similar patterns of presence/absence of CyCOG products within each genus (Figure 4), and when subjected to *k*-means analysis, these patterns represent the same major groups identified in the AAI (Figure 2) and phylogenomic (Figure 3) analyses. Grouping into *k*-means is show in Table 2. When *k* = 2, the division is broad, between the Marine groups (including the Symbiont *Synechospongium*) and Freshwater/thermal. When *k* is raised to 3, the division is between Marine, Symbiont and Freshwater/thermal. When *k* = 4, the division is between Marine, Symbiont, Freshwater and Thermal genera. For each respective *k* value, the data shows that: i) The broadest ecogenomic divide is between genomes of marine and freshwater/thermal environments; ii) the Symbiont group is then separated, suggesting that its symbiotic lifestyle has led to a different pattern of CyCOG presence/absence within the Marine group (Slaby & Hentschel, 2017) and iii) Within the Freshwater/thermal group, the Freshwater and Thermal group display distinct patterns. There was little difference within genera of the Marine/oceanic and Marine/coastal groups. This was perhaps surprisingly, as some genomes from these groups come from very different environments, such as the *Regnicoccus* genome which are sourced from both temperate estuarine waters (the type species WH 5701 was isolated from the Long Island Sound, USA) (Fuller et al. 2003) and extreme environments such as the Ace Lake, in the Vestfold Hills of Antarctica (strain SynAce01) (Powell et al. 2005). The new genus *Lacustricoccus* is also surprisingly grouped within the Marine/coastal group, as genomes from this genus were sourced from brackish water reservoirs (Cabello-Yevez et al. 2017, 2018).

## CONCLUSION

It is timely to establish a genome-based taxonomy for SC (Gevers et al. 2005, Stackebrandt 2006). With the advent of next generation sequencing and increasingly available sequence data, there has been a transition from the former paradigm of a ‘polyphasic’ taxonomy towards a genomic taxonomy (Thompson et al. 2015). Examining prokaryotic taxonomy using the organisms’ whole genome would be able to capture meaningful relationships and define monophyletic groups, capturing their rate of evolution across taxonomic ranks (Hugenholtz et al. 2016, Parks et al. 2018). In their large-scale analysis, Parks and colleagues (2018) examined over 18000 genomes and divide the *Synechococcus* in at least 5 genera, but, these authors do not delve further into the detailed taxonomic analyses of the taxon. To the best of our knowledge, there is not a consensus on whether the *Synechococcus* form a monophyletic clade. This may be the case for specific marine or freshwater lineages, but when examined in the context of the *Cyanobacteria* phylum, the genus as presently classified is paraphyletic or polyphyletic as demonstrated here (Walter et al. 2017). Our advanced genomic taxonomy analyses demonstrate the heterogeneous nature of the SC collective. This study brings new insights into the taxonomic structure of SC collective with the evident distinction of 11 genera. We anticipate that this newly proposed taxonomic structure will be useful for further environmental surveys and ecological studies (Arevalo et al. 2019), including those targeting the identification of populations, ecotypes and species.

## ACKNOWLEDGEMENTS

The authors thank CAPES and CNPq for funding.

